# Biomechanical comparison of plate materials and designs for subcondylar fracture fixation: An *in silico* assessment

**DOI:** 10.1101/2023.08.07.552268

**Authors:** Anoushka Gupta, Rajdeep Ghosh, Kaushik Dutta, Abir Dutta, Kaushik Mukherjee

## Abstract

Mandibular subcondylar fractures present challenges in fixation due to high complication rates and the uncertainty in selecting optimal plate design and material. Previous computational studies have primarily relied on simplified models, point loading, and plate-only analyses. This study examines the combined influence of fixation plate design and material properties on fracture stability, with interfragmentary displacement as a key indicator of post-surgical stability. A finite element approach implemented in ANSYS Mechanical APDL v2022 R2, incorporating soft tissues such as the periodontal ligament, was employed to assess fixation stability using five materials – Nitinol, Magnesium alloys, Titanium alloys (Ti-6Al-4V and Ti-29Nb-13Ta-4.6Zr), and Stainless Steel 316L – across four plate designs under ipsilateral and contralateral molar clenching simulated using physiologically-mimetic muscle forces. Results showed a reduction in mandibular strain (up to 4% from around 1635 µ*ε* to 1569 µ*ε*) with increase in the plate stiffness during ipsilateral clenching, while for contralateral clenching, this trend (up to 5.4% from around 1482 µ*ε* to 1402 µ*ε*) was observed only for double mini and lambda plates due to enhanced resistance to bending moments from additional screw placement. Among designs, the double mini plate demonstrated the greatest reduction in the interfragmentary gap (by 77% during ipsilateral clenching; by 58% during contralateral clenching) and emerged as the most stable option across materials. Regarding materials, Titanium alloys (TNTZ and Ti-6Al-4V) were mechanically preferable, demonstrating higher safety margins based on factory of safety (FoS) ratios (TNTZ: FoS 5.57 ipsilateral, 3.21 contralateral; Ti-6Al-4V: FoS 5.05 ipsilateral, 2.98 contralateral), indicating reduced risk of yielding under functional loading. These findings underscore the critical interplay between screw configuration, material selection, and loading conditions, offering valuable guidance for implant design optimization.

## 1. Introduction

The subcondylar region of the mandible is one of the most frequently fractured areas of the face [1,2], often due to vehicular accidents or interpersonal violence. Presently, Open Reduction and Internal Fixation (ORIF) is the standard treatment for mandible fractures [3]. However, ORIF has a complication rate of 10-30%, which often demands secondary surgeries and adds to financial burdens [4]. The narrow anatomy and unique physiology of subcondylar fractures make fixation particularly challenging. The choice of fixation plate material and geometric configuration is critical for a successful surgery. For optimal patient outcomes, the implant’s stiffness should match with that of a native bone tissue, and the geometric configuration should facilitate effective load transfer across the narrow subcondylar region. Recently, three-dimensional plates such as trapezoid, strut, and lambda plates have been specifically designed for subcondylar fractures [5]. Additionally, the use of double mini plates has gained support for providing stability in fixation [6].Although there are different types of clinically approved materials, the Ti-6Al-4V alloy is considered the gold standard for its favourable biomechanical strength and biocompatibility. However, Ti-6Al-4V has higher elastic modulus as compared to the cortical bone [7], which causes stress shielding, resulting in an implant-removal rate of 10-12% [8]. Alternatively, steel (AISI SS 316L), has been popularly used for its affordability. However, steel has two times higher density (8.1-8.9 g/cm3) than titanium (4.4-4.5 g/cm3) along with higher modulus of 210 GPa [8,9]. In addition, these common metallic alloys are prone to corrosion, such as pitting, crevice corrosion and corrosion [10]. To overcome these issues, new alloys are being developed by including non-cytotoxic elements such as Niobium (Nb), Zirconia (Zr) and Tantalum (Ta) to reduce modulus and impart anti-corrosion ability. One such alloy which has been used for fracture fixation plate is Ti-29Nb-13Ta-4.6Zr (aged) with bio-inertness [11] and lower modulus of ∼80 GPa [12] (as compared to 110 GPa of Ti-6Al-4V [13]). The reduced elastic modulus of this alloy brings its stiffness closer to that of cortical bone, which can enhance load sharing between the fixation plate and mandibular bone. Improved load sharing promotes more physiological strain distribution and may mitigate stress shielding effects during healing in mandibular fixation, while maintaining adequate mechanical stability and corrosion resistance [14,15]

The alternative materials, such as Mg WE43 alloy (Mg-3.5Y-2.3Nd-0.5Zr), have a lower elastic modulus (44.2 GPa [16]) favouring neo-bone growth and integration [8]. Further advantages of Mg include low densities (1.74-2.0 g/cm^3^ [8]) and abundant presence of Mg^2+^ cation in the body [7]. Mg WE43 alloy has an appropriate biodegradation rate for preventing non-union [17]. Another popular material being used in clinics for orthodontic braces is solid nitinol (NiTi), owing to its excellent biocompatibility, mechanical properties, and resistance to corrosion [18]. Furthermore, nitinol has less modulus (38.5 GPa [19]), which makes it a suitable choice for mandibular fracture fixation plates [20].

The finite element analysis (FEA) technique has been proven to be a preferred method to assess the feasibility of mandibular fixation methods [21,22]. FEA for mandibular fractures has been validated using 3D-printed polymer mandibles [23] and has been widely applied to investigate different scenarios with various plate designs, plate thicknesses, materials and bone conditions such as atrophy [22, 24–27], as also highlighted in a recent review article [28]. For instance, Orassi et al. [16] compared Ti-6Al-4V with Mg WE43 for condylar neck fractures using a double mini-plate configuration across plate thicknesses and found that Mg WE43 achieved strength comparable to titanium while offering the advantages of biodegradability. Prasadh et al. [29] also compared the same materials for mandibular angle fracture, and Jung et al. [30] compared four materials, but with very simplified models. Hassani et al. [24] reported that the location of screws played a crucial role in subcondylar fracture fixation. Despite these advances, many of these studies assigned simplified, homogeneous material properties rather than heterogeneous cortical–cancellous definitions; used idealized loading; and omitted the periodontal ligament (PDL) and temporomandibular joint fibrocartilage. Moreover, a systematic evaluation of fracture-site micromotion arising from the combined effects of plate geometry and material stiffness under physiologically realistic mandibular loading remains lacking. To address these gaps, we developed a high-fidelity FE mandible model that explicitly represents teeth, the PDL, condylar fibrous tissues, and distinct cancellous/cortical bone, and we simulated a physiologically realistic mastication cycle [31]. In addition, the influences of the chosen five different materials, including nitinol and less rigid Ti alloy, as well as different designs, have not been explored yet. To enable a focused evaluation of material–design interactions, four plate configurations with favourable biomechanical performance, reported in our earlier study [31], were selected for detailed analysis.

The primary aim of this study is to investigate how material stiffness modulates the fixation stability across clinically relevant plate designs in subcondylar mandibular fractures, by focussing on the interfragmentary displacement as a mechanistic indicator of the initial timepoints immediately following the surgical fixation. Specifically, in this study, a comparative biomechanical evaluation of subcondylar fixation has been carried out for reconstructed mandibles with the following combinations of five different implant materials, namely Nitinol, Magnesium Alloy WE43, a low rigidity Titanium Alloy (TNTZ: Ti–29Nb–13Ta–4.6Zr), a common Titanium Alloy (Ti-6Al-4V) and Stainless Steel (SS 316L) and four implant designs, namely trapezoid, strut, lambda and double mini plates.

## 2. Materials and Methods

Anonymized computed tomography (CT) images with a pixel resolution of 512×512, pixel size of 0.439 mm and slice thickness of 0.75 mm of a healthy subject were obtained after necessary approval from the ethics committee of Guru Nanak Institute of Dental Sciences and Research, Kolkata, India, for the development of the intact mandible model. A virtual fixation with four plates was performed on the modeled fractured mandible. Thereafter, the plates were assigned with five different effective material properties, to investigate the influence of the materials on the load-transfer across the mandible.

### 2.1. Development of 3D model of intact mandible

The three-dimensional mandible model (Figure 1a) was developed from an anonymized CT-scan of a 20-year-old healthy male, with consent obtained at Guru Nanak Institute of Dental Sciences and Research, India. The mandible was segmented into cortical and cancellous bone, teeth, 0.2 mm thick periodontal ligament (PDL), and 0.3mm thick condylar fibrocartilage using Mimics 24.0 (Materialise, Leuven, Belgium) [31–35]. Temporal bone was modelled as rigid block with concave depression [31–34] as shown in Figure 1. A global mesh convergence study was performed, and mesh sizes were chosen for the model having <1% deviation in the state of maximum tensile strain. Based on the mesh convergence study, an element size of 0.2 – 1 mm was chosen for discretization. The entire model was discretized with 10-noded tetrahedral elements.

**Figure 1.**
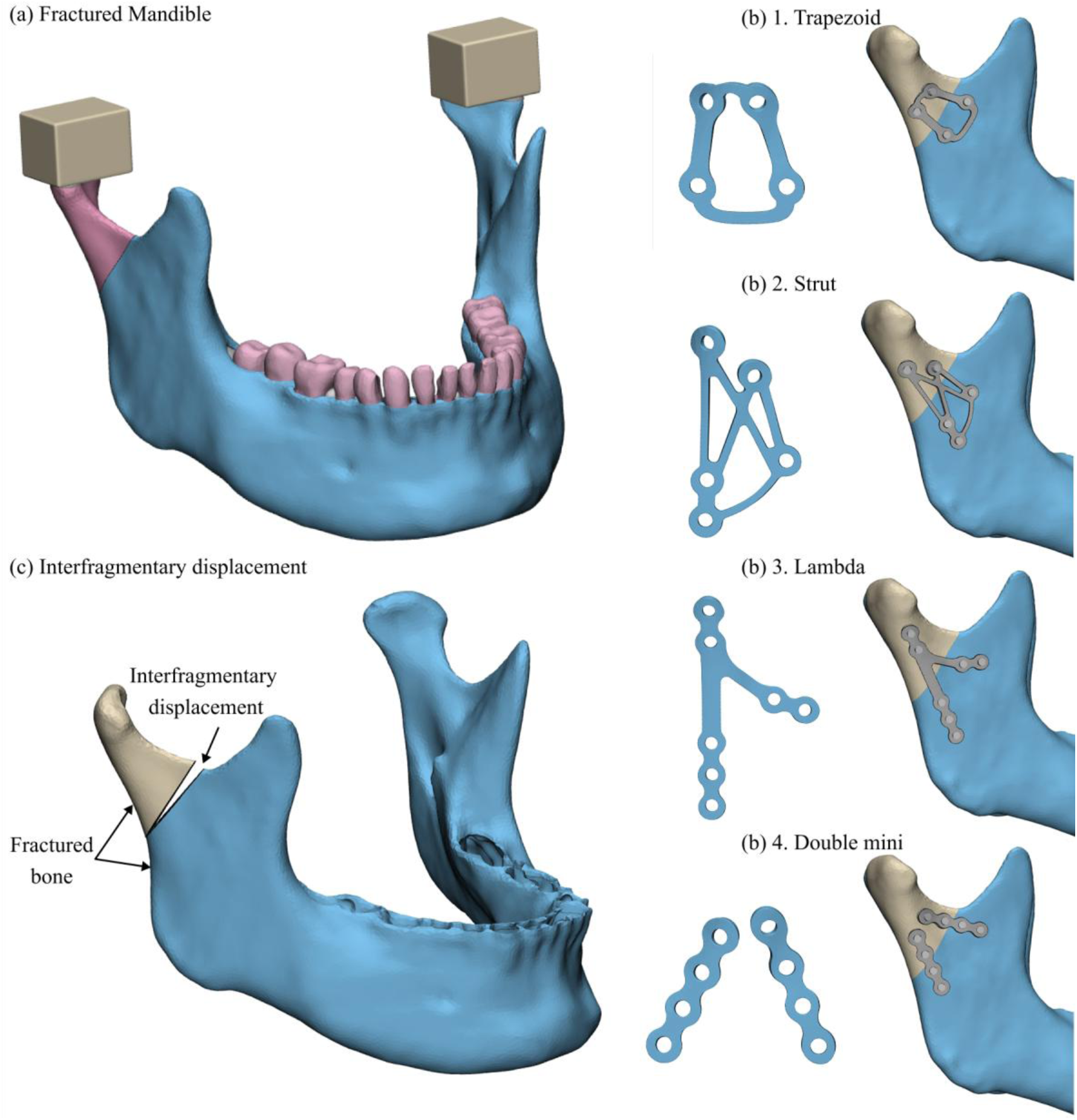
(a) Developed model of fractured mandible (b) Developed plate models and corresponding mandibular reconstructions: 1. Trapezoid, 2. Strut, 3. Lambda, 4. Double mini plates, (c) Interfragmentary displacement calculated at the maximum opening end of the osteotomy site

### 2.2. Development of implanted mandible

A standardized osteotomy line [36] for subcondylar fracture, from the mid-point of the ramus to the bottommost point of sigmoid notch, was simulated in 3-Matic 16.0 (Materialise, Leuven, Belgium). Four different plate designs (trapezoid, strut, lambda and double mini) (Figure 1b) were chosen. The patient-specific plate designs were inspired from the commercially available plate designs (DePuy Synthes, PA). The screws were modelled as cylinders with a diameter of 2 mm and a length of 6 mm [37, 38]. To model the cold-welded locking screws, no relative movement was allowed between the screw and the plate. The double mini plates were fixed in an angulated manner [36] and the trapezoid, strut and lambda plates were fixed based on the recommended guidelines[5]. All the reconstructed mandibles were meshed using 10-noded tetrahedral elements using the same FE model development procedure as described in section 2.1 and had a similar number of elements (∼14,00,000).

### 2.3. Assignment of Material Properties

Cancellous bone was assigned with voxel-based non-homogeneous isotropic properties. For each element, the apparent density (⍴) was calculated corresponding to its gray-scale value (in HU) following the relationship [31]:

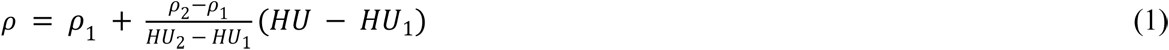

where (*ρ*_1_, *HU*_1_) represented the no-bone condition (0.022 g/cm^3^, 0 HU) and (*ρ*_2_, *HU*_2_) represented the hard-bone condition (1.73 g/cm^3^, 2600 HU). The negative densities corresponding to <0 HU were flushed to the minimum values corresponding to the no-bone condition [39–41]. A power law relationship was used to calculate the element-specific Young’s modulus (*E*) of cancellous bone based on its apparent density [31,42]:

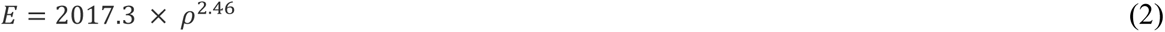

Cortical bone was assigned with region-specific orthotropic properties based on the literature [13,31]. The rest of the structures, including the plates and screws were modelled as homogenous linear isotropic [13,31]. The teeth were further divided into enamel and dentin with different isotropic material properties [13,31]. Table 1 provides a summary of the material properties assigned to each component. The plates were assigned with five different material properties as presented in Table 2 [9,12,13,16,19]. Titanium alloy (Ti-6Al-4V) properties (E = 110 GPa, ν = 0.3) [13] were assigned to the screws in all the cases.

**Table 1.** Material properties of components of Finite Element model.

| Component | $E_X$<br>(GPa) | $E_Y$<br>(GPa) | $E_Z$<br>(GPa) | $\nu_{XY}$ | $\nu_{YZ}$ | $\nu_{XZ}$ | $G_{XY}$<br>(MPa) | $G_{YZ}$<br>(MPa) | $G_{XZ}$<br>(MPa) |
| --- | --- | --- | --- | --- | --- | --- | --- | --- | --- |
| Symphysis and mental foramen | 23 | 10 | 15 | 0.3 | 0.3 | 0.3 | 4800 | 3600 | 6200 |
| Gonial angle | 20 | 11 | 12 | 0.3 | 0.3 | 0.3 | 4800 | 5300 | 6000 |
| Rest of cortical | 17 | 6.9 | 8.2 | 0.3 | 0.3 | 0.3 | 2800 | 2900 | 4600 |
| Cancellous | $1 \times 10^{-4} - 4.33$ | - | - | 0.3 | - | - | - | - | - |
| Enamel | 80 | - | - | 0.3 | - | - | - | - | - |
| Dentin | 17.6 | - | - | 0.25 | - | - | - | - | - |
| Condylar Fibrous Tissue | 0.006 | - | - | 0.25 | - | - | - | - | - |
| Periodontal ligament | 0.003 | - | - | 0.45 | - | - | - | - | - |
| Blocks | 50000 | - | - | 0.3 | - | - | - | - | - |
| Screws (Ti-6Al-4V) | 110 |  |  | 0.3 |  |  |  |  |  |

**Table 2.** Material properties of plates.

| **Material** | ***Elastic modulus***<br>*($E_X$ in GPa)* | **Poisson's ratio**<br>($\nu$) | ***Yield Strength***<br>*(MPa)* | ***Reference*** |
| --- | --- | --- | --- | --- |
| Nitinol | 38.5 | 0.3 | 200 | Moghaddam et al. 2016 [17] |
| Mg WE43 | 44.2 | 0.27 | 162 | Orassi et al. 2022 [14] |
| TNTZ | 80 | 0.3 | 864 | Niinomi et al. 1998 [12] |
| Ti-6Al-4V | 110 | 0.3 | 900 | Dutta et al. 2019 [13] |
| SS 316L | 210 | 0.3 | 380 | Xie et al. 2013 [9] |

### 2.4. Applied loading and boundary conditions

All the intact and reconstructed mandibles were simulated under the contralateral left unilateral molar clenching (LMOL) and the ipsilateral right unilateral molar clenching (RMOL) to mimic ‘alternating bilateral clenching pattern’ [43,44]. A total of seven muscles were considered in the study: superficial and deep masseter muscles (SM and DM), medial and inferior lateral pterygoid (MP and ILP), and anterior, middle, and posterior temporalis (AT, MT and PT) [13] The magnitude of the muscle forces was distributed over the nodes representative of the muscle attachment sites [13]. The top surfaces of the rigid temporal blocks were fixed during the clenching tasks [13]. The details of the applied muscle forces and boundary conditions are shown in Table 3.

**Table 3.**
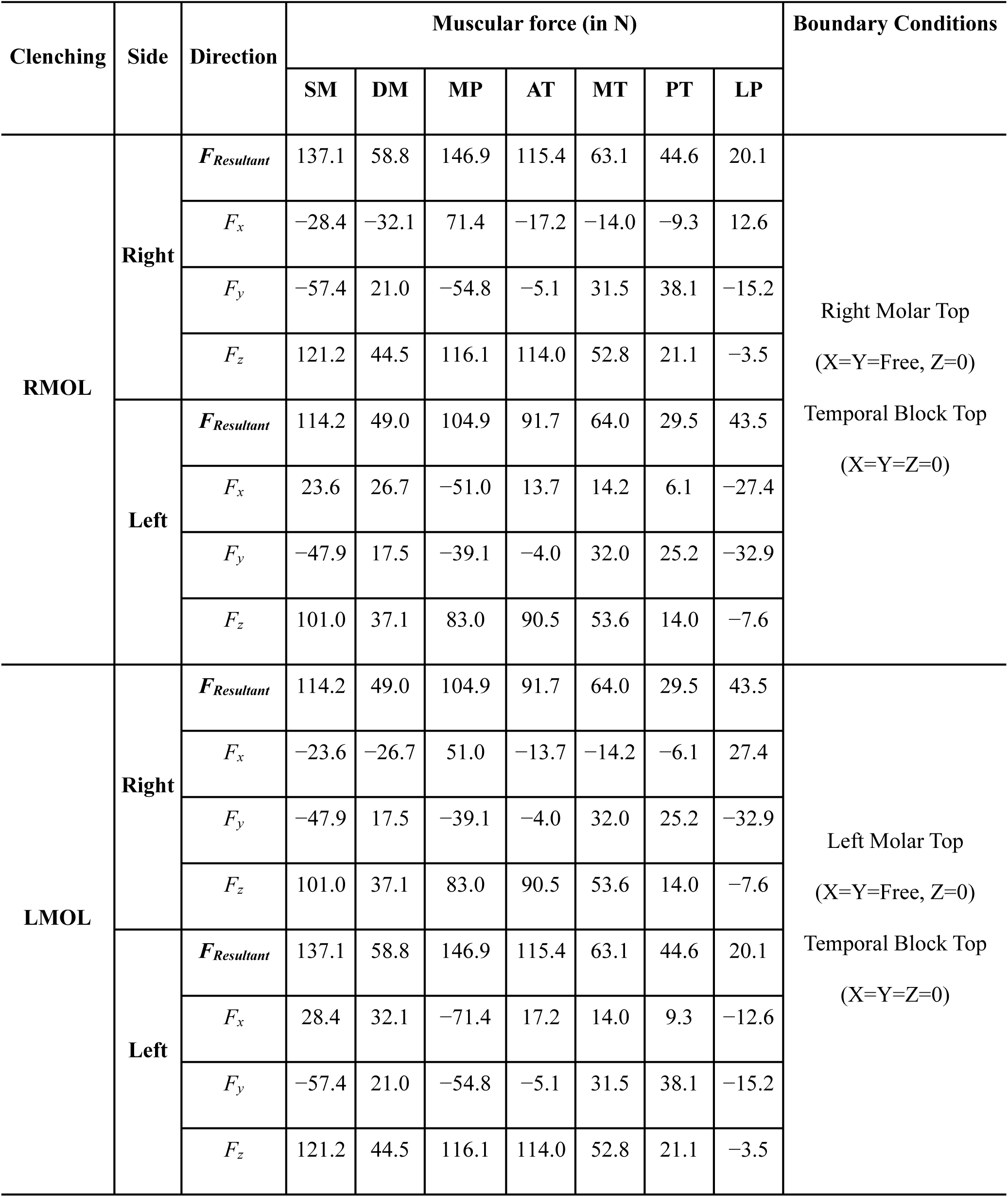
Muscle Forces and Boundary Conditions Applied.

A standard frictional contact pair was defined with a frictional coefficient of 0.4 [31,45,46] at the plate-cortical and fracture interfaces. Rest of the surfaces between the adjacent parts, such as the screw with the plate, cancellous and cortical, were modelled as bonded. While this assumption of bonded interfaces facilitates analysis of global biomechanical behaviour, it does not permit prediction of micro-movements or loosening at the interfaces, which should be considered when interpreting results. An augmented Lagrangian was employed in the FE study.

### 2.5. Verification and validation of the FE model generation method

As the CT scan was from a living subject, direct one-to-one experimental validation of the FE model was not possible. Nevertheless, the maximum principal strains and stresses in the intact mandible during right molar clenching were found to be of quantitatively similar magnitude (stress: 15MPa; strain: ∼1200 με, Supplementary Figure S1) at qualitatively similar regions (for example, high stresses near the anterior buccal body and symphysis region of the mandible) as reported in literature [33]. Moreover, the reaction force generated at the constrained right molar teeth was ∼850 N for the intact mandible during the right unilateral molar clenching task. In the present model, this reaction force represents the resultant occlusal bite force produced by the applied muscle forces under equilibrium conditions and is within the reported range of occlusal forces for healthy humans (365–1230 N) [47]. Also, the direction of the overall deformation was found to be towards the working (chewing) side (Supplementary Figure S1), as found in earlier reports [33,48]. Hence, the reliability and accuracy of the FE modelling methodology were verified and adopted for all the reconstructed mandibles.

### 2.6. Interpretation of results

The maximum principal strains and stresses in the reconstructed mandible were estimated to evaluate the efficacy of devices and risk of non-union and secondary fractures. The von Mises stresses of the plates were calculated to predict the failure of ductile plate materials. 95th percentile values of all the stresses and strains were computed over the entire structure (bone or implant and screws) to exclude singularities around the constrained regions [33].In addition, the interfragmentary displacement, defined as the distance between the two bone surfaces on both sides of the osteotomy line (Figure 1c), was also computed to understand the fixation stability.

## 3. Results

The principal stresses, strains and von Mises stresses in the cortical bone and implants, respectively, are presented below. Further, different virtual fractures created at the subcondylar site due to influence of chewing laterality, fixation design and material are discussed. Only the von Mises stress distributions in the plates have been presented in detail, whereas the results for the screws are presented in the supplementary data.

### 3.1. Principal stresses and strains in the mandible

The maximum principal stresses in the intact mandible were found on the buccal border below the right or left molar teeth under RMOL (14.8 MPa) or LMOL (14.1 MPa), respectively (Supplementary Figure S1). The maximum principal stresses in all the reconstructed mandibles (15-18 MPa) were higher as compared to the intact mandible. However, the different material properties of plates resulted in minimal differences (± 0.2-0.5 MPa) in maximum principal stresses in the reconstructed mandibles (Supplementary Table S1). Among different plate designs, highest maximum principal stresses (∼18 MPa under RMOL, ∼16 MPa under LMOL) were observed in the mandible reconstructed with trapezoid plate. Whereas the lowest principal stresses were found in case of lambda plate (∼17 MPa) under RMOL and strut plate (∼15 MPa) under LMOL (Supplementary Table S1).

The maximum principal strains (1288 με under RMOL and 1240 με under LMOL) occurred along the anterior border near the condyle and coronoid of intact mandible (Supplementary Figure S3). In reconstructed mandibles, there were higher strain values (1383 - 1635 με) than those observed in the intact mandible (Table 4). For different materials, an increase in the young’s modulus of the plates resulted in an increase of its stiffness which was also associated with a reduction in the maximum principal strains (Table 4). Only strut and trapezoid plates exhibited an opposite trend of increasing strains (∼20 – 100µε) in reconstructed mandibles under LMOL loading condition with an increase in plate stiffness (Table 4). Among all plate designs, the strut plate resulted in the least maximum principal strain in reconstructed mandibles for most of the materials (Table 4).

**Table 4.** Maximum principal strains (µε) for reconstructed mandibles during RMOL and LMOL for different implant materials.

| Loading conditions | Implant type | Nitinol | Mg WE43 | TNTZ | Ti-6Al-4V | SS 316L |
| --- | --- | --- | --- | --- | --- | --- |
| **RMOL** | Trapezoid | 1605 | 1609 | 1614 | 1611 | 1595 |
|  | Strut | 1541 | 1538 | 1528 | 1522 | 1517 |
|  | Lambda | 1572 | 1568 | 1554 | 1549 | 1543 |
|  | Double mini | 1635 | 1628 | 1603 | 1558 | 1569 |
| **LMOL** | Trapezoid | 1478 | 1481 | 1496 | 1498 | 1495 |
|  | Strut | 1383 | 1385 | 1411 | 1429 | 1466 |
|  | Lambda | 1482 | 1476 | 1451 | 1434 | 1402 |
|  | Double Mini | 1511 | 1506 | 1484 | 1473 | 1461 |

### 3.2. Stress and strains in implant components

Maximum von Mises stresses in the plate were found to increase with an increase in plate material modulus (Table 5). Under RMOL, peak plate stresses ranged from 111 MPa (Double mini, nitinol) to 456 MPa (Lambda, SS316L). Under LMOL, stresses increased substantially, ranging from 171 MPa (Lambda, nitinol) to 520 MPa (Trapezoid, SS316L). Among the plate designs, the lambda and trapezoid plates generally exhibited higher peak stresses compared to the strut and double mini plates under both loading conditions. The double mini plate consistently showed the lowest stresses across materials, particularly under RMOL. For titanium (both Ti-6Al-4V and TNTZ) fixation plates, the maximum von Mises stresses did not exceed the yield strength for all designs (Table 5). Notably, TNTZ plates demonstrated lower stress-to-yield ratios compared to Ti-6Al-4V despite a lower elastic modulus, indicating improved relative safety margins. For nitinol, only the strut plate exhibited maximum von Mises stress lower than its yield strength (Table 5). Whereas for other two materials (Mg WE43 and SS316L), all plates experienced maximum von Mises stresses beyond corresponding yield value, indicating reduced safety margins (Table 5). For a concise quantitative comparison, the factor of safety (FoS), defined as the ratio of material yield strength to maximum von Mises stress, was evaluated for the double mini plate configuration as a representative case. Under RMOL, the FoS values were approximately 1.80 (Nitinol), 1.36 (Mg WE43), 5.57 (TNTZ), 5.05 (Ti-6Al-4V), and 1.58 (SS 316L). Under LMOL, the FoS decreased to approximately 0.97 (Nitinol), 0.75 (Mg WE43), 3.21 (TNTZ), 2.98 (Ti-6Al-4V), and 0.99 (SS 316L). This comparison demonstrates the substantially higher safety margins provided by the titanium alloys, whereas Mg WE43, SS 316L, and Nitinol approach or fall below unity under contralateral loading, indicating an increased likelihood of yielding. A marginal increase in von Mises stresses was found in the posterior part of the plates under RMOL as compared to those under LMOL (Figures 2 - 3). It was observed that there is a higher stress distribution in the anterior arm of the trapezoid plate under LMOL (Figure 3).

**Figure 2.**
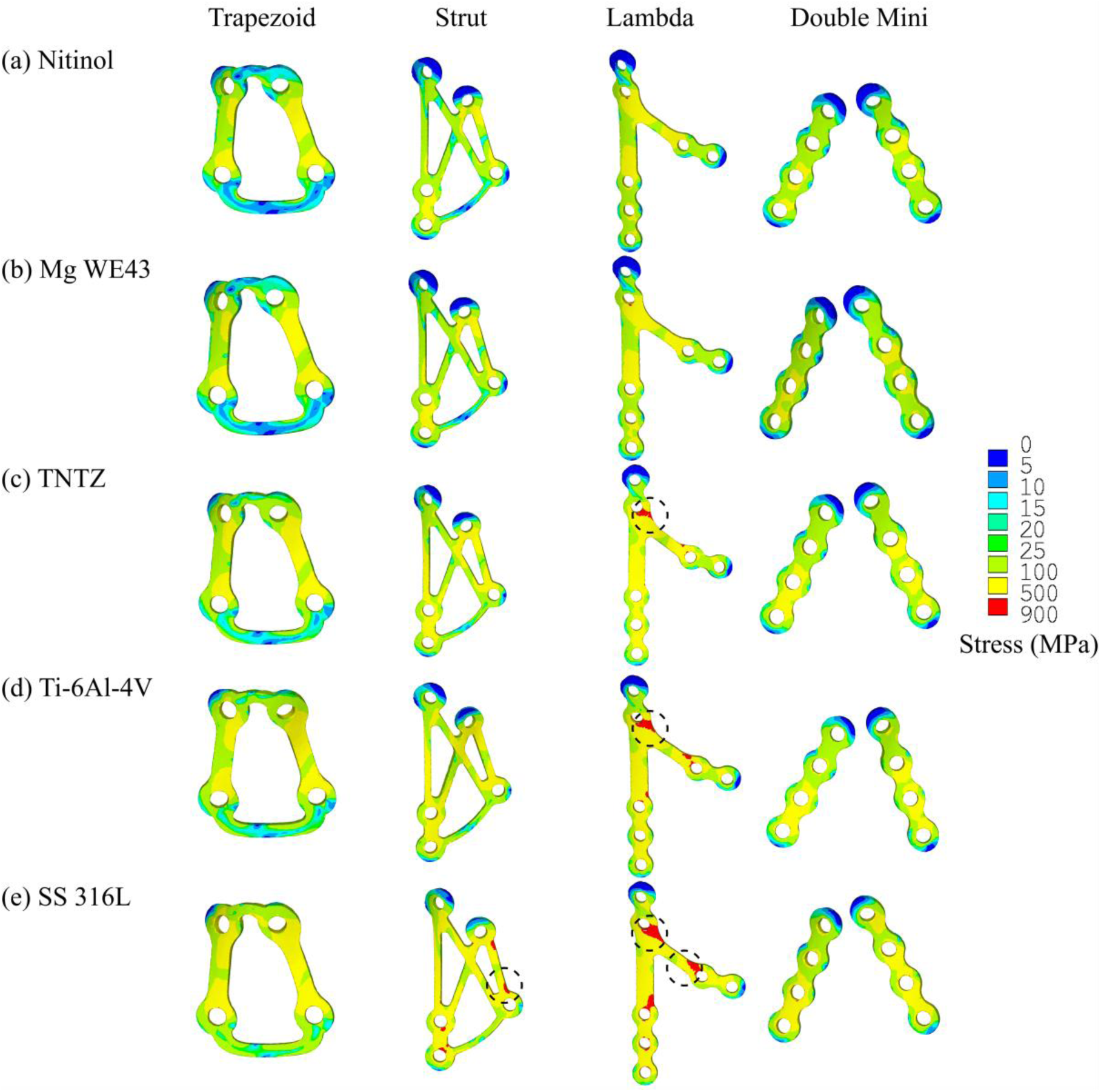
Von Mises stress distribution in trapezoid, strut, lambda and double mini plates for (a) Nitinol, (b) Mg WE43, (c) TNTZ, (d) Ti-6Al-4V and (e) SS 316L under RMOL

**Figure 3.**
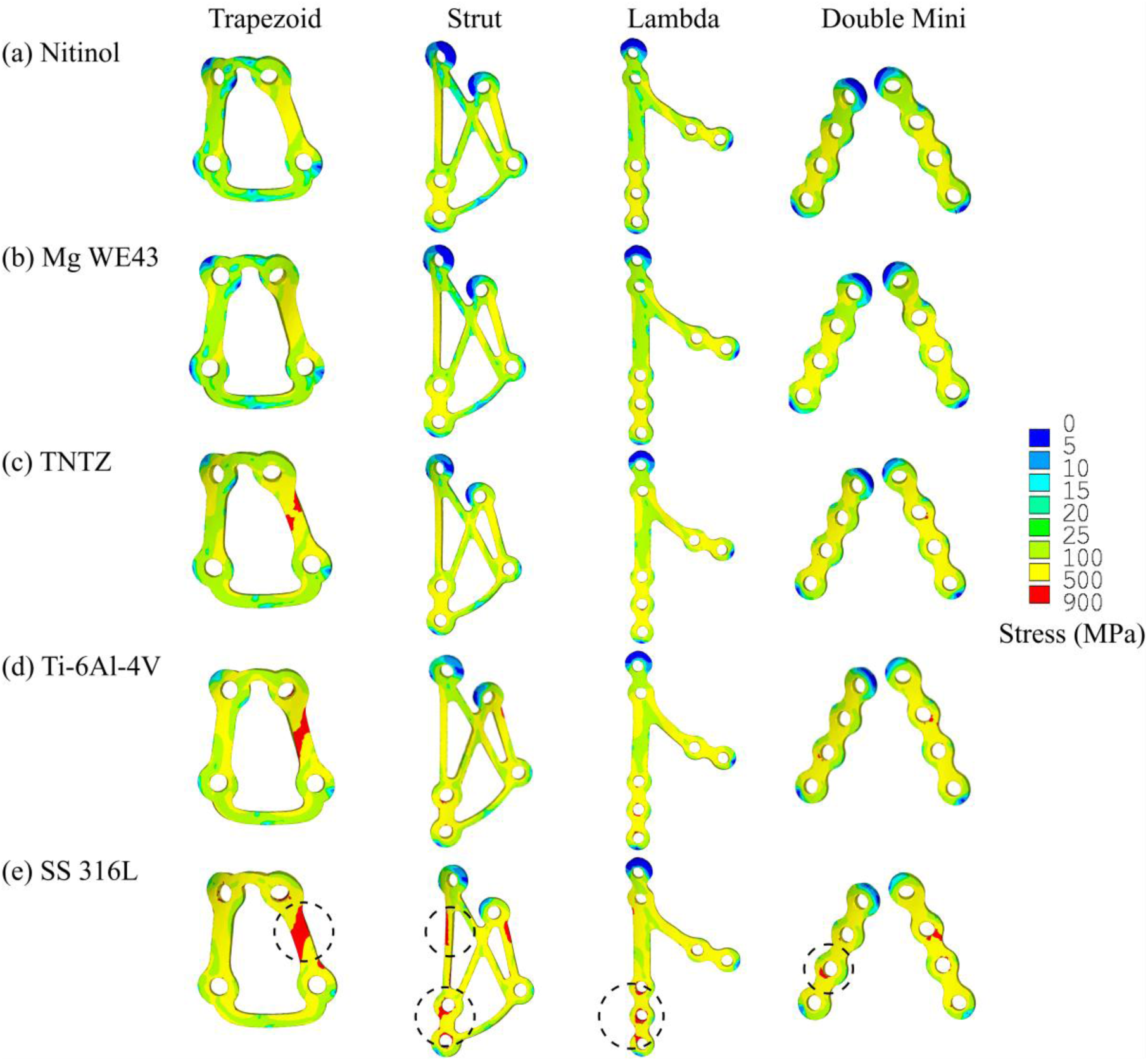
Von Mises stress distribution in trapezoid, strut, lambda and double mini plates for (a) Nitinol, (b) Mg WE43, (c) TNTZ, (d) Ti-6Al-4V and (e) SS 316L under LMOL

**Table 5.**
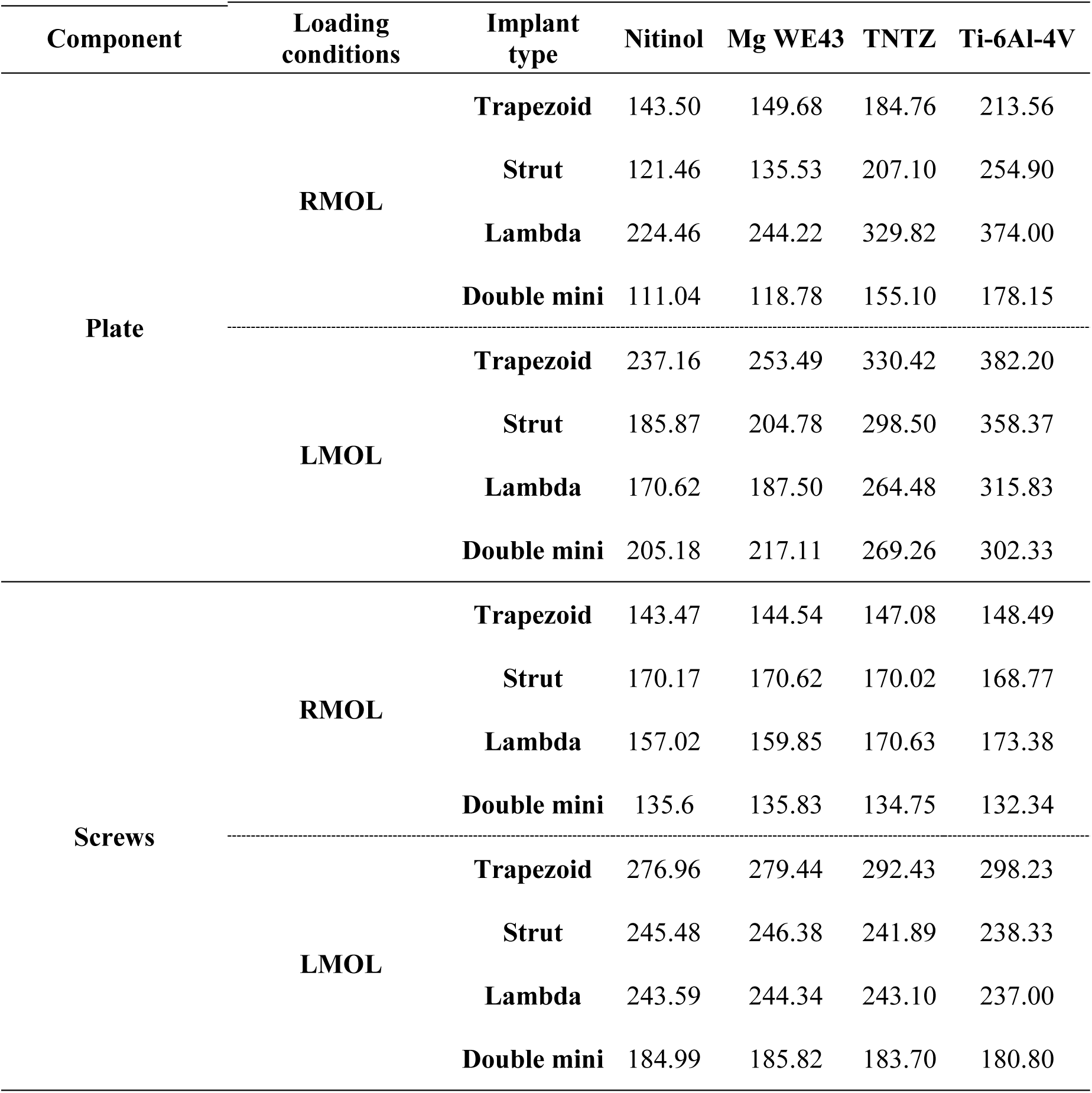
Maximum von Mises stresses (MPa) in plates and screws during RMOL and LMOL for different implant materials.

| Component | Loading conditions | Implant type | Nitinol | Mg WE43 | TNTZ | Ti-6Al-4V |
| --- | --- | --- | --- | --- | --- | --- |
| Plate | RMOL | Trapezoid | 143.50 | 149.68 | 184.76 | 213.56 |
|  |  | Strut | 121.46 | 135.53 | 207.10 | 254.90 |
|  |  | Lambda | 224.46 | 244.22 | 329.82 | 374.00 |
|  |  | Double mini | 111.04 | 118.78 | 155.10 | 178.15 |
|  | LMOL | Trapezoid | 237.16 | 253.49 | 330.42 | 382.20 |
|  |  | Strut | 185.87 | 204.78 | 298.50 | 358.37 |
|  |  | Lambda | 170.62 | 187.50 | 264.48 | 315.83 |
|  |  | Double mini | 205.18 | 217.11 | 269.26 | 302.33 |
| Screws | RMOL | Trapezoid | 143.47 | 144.54 | 147.08 | 148.49 |
|  |  | Strut | 170.17 | 170.62 | 170.02 | 168.77 |
|  |  | Lambda | 157.02 | 159.85 | 170.63 | 173.38 |
|  |  | Double mini | 135.6 | 135.83 | 134.75 | 132.34 |
|  | LMOL | Trapezoid | 276.96 | 279.44 | 292.43 | 298.23 |
|  |  | Strut | 245.48 | 246.38 | 241.89 | 238.33 |
|  |  | Lambda | 243.59 | 244.34 | 243.10 | 237.00 |
|  |  | Double mini | 184.99 | 185.82 | 183.70 | 180.80 |

Maximum principal strains in the plates decreased consistently with increasing elastic modulus under both loading conditions (Table 6). Across materials, the strain reduction from nitinol to SS316L was substantial. For example, under LMOL in the trapezoid plate, strain decreased from 6080 µε (nitinol) to 2277 µε (SS316L), corresponding to approximately a 63% reduction. A similar decreasing trend with increasing stiffness was observed for all implant types. For all implant designs, nitinol exhibited the highest strains, followed by Mg WE43, TNTZ, Ti-6Al-4V, and SS316L. Under RMOL, maximum principal strains ranged from 928 µε (Double mini, SS316L) to 4180 µε (Lambda, nitinol). Under LMOL, strains increased substantially, ranging from 1493 µε (Double mini, SS316L) to 6080 µε (Trapezoid, nitinol).

**Table 6.** Maximum principal strains (µε) in plates under RMOL and LMOL for different implant materials.

| **Loading conditions** | **Implant type** | **Nitinol** | **Mg WE43** | **TNTZ** | **Ti-6Al-4V** | **SS 316L** |
| --- | --- | --- | --- | --- | --- | --- |
| **RMOL** | **Trapezoid** | 3240 | 2941 | 2022 | 1693 | 1221 |
|  | **Strut** | 2494 | 2398 | 2016 | 1784 | 1364 |
|  | **Lambda** | 4180 | 3952 | 3014 | 2465 | 1638 |
|  | **Double Mini** | 2383 | 2210 | 1603 | 1329 | 928 |
| **LMOL** | **Trapezoid** | 6080 | 5642 | 3992 | 3301 | 2277 |
|  | **Strut** | 4066 | 3802 | 2891 | 2474 | 1789 |
|  | **Lambda** | 3554 | 3356 | 2653 | 2310 | 1738 |
|  | **Double Mini** | 4452 | 4087 | 2845 | 2325 | 1493 |

Both the stresses and strains in plates were generally higher under LMOL than RMOL (Tables 5 – 6). For example, in the trapezoid plate, stress increased from 214 MPa (Ti-6Al-4V, RMOL) to 382 MPa (LMOL), representing approximately a 79% increase. A similar trend was observed for SS316L plates, where stress increased from 300 MPa to 520 MPa (∼73% increase). Moreover, strain increased from 3240 µε (Nitinol, RMOL) to 6080 µε (LMOL), indicating a 88% increase in strain. Similar observations were made for other plates and materials as well.

Overall, the maximum von Mises stresses in the screws were not influenced by plate materials to a large extent (Table 5, Supplementary Figures S1 – S2). Under RMOL, screw stresses ranged from 126 MPa to 173 MPa. Under LMOL, stresses increased substantially, ranging from 170 MPa to 304 MPa. LMOL resulted in approximately 60–100% higher screw stresses compared to RMOL, depending on plate design. For instance, trapezoid screw stresses increased from approximately 148 MPa (RMOL, Ti-6Al-4V) to 298 MPa (LMOL), corresponding to a ∼101% increase. Despite this increase, screw stresses remained below the yield strength of titanium in all configurations, indicating adequate immediate mechanical safety (Table 5, Supplementary Figures S2 – S3). The double mini configuration consistently exhibited the lowest screw stresses under both loading conditions, whereas trapezoid and strut designs showed comparatively higher screw loading under LMOL.

### 3.3. Interfragmentary displacement

For all plate designs and load cases, the interfragmentary displacements were found to be decreasing with increase in stiffness of fixation plates (Table 7). Among all plate designs, the least interfragmentary displacement (6 – 44.90 µm) was found with the double mini plate, whereas the highest interfragmentary displacement (86.80 – 530 µm) was found with lambda plate. For lambda plate, interfragmentary displacement was higher under RMOL as compared to LMOL, whereas, for double mini plate, interfragmentary displacement was higher under LMOL as compared to RMOL (Table 7). For trapezoid and strut plates, the interfragmentary displacements for nitinol and MgWE43 were marginally higher under RMOL than LMOL (Table 7). However, for TNTZ, Ti-6Al-4V and SS316L, trapezoid and strut plates exhibited higher interfragmentary displacements under LMOL than RMOL (Table 7).

**Table 7.** Interfragmentary displacement (µm) during RMOL and LMOL for different implant materials.

| **Loading conditions** | **Implant type** | **Nitinol** | **Mg WE43** | **TNTZ** | **Ti-6Al-4V** | **SS 316L** |
| --- | --- | --- | --- | --- | --- | --- |
| **RMOL** | **Trapezoid** | 138.57 | 126.00 | 86.91 | 73.00 | 55.40 |
|  | **Strut** | 131.00 | 128.00 | 111.56 | 90.00 | 70.30 |
|  | **Lambda** | 530.00 | 507.00 | 383.00 | 300.00 | 186.00 |
|  | **Double Mini** | 27.00 | 24.60 | 15.18 | 11.00 | 6.10 |
| **LMOL** | **Trapezoid** | 116.90 | 112.60 | 92.56 | 80.00 | 68.40 |
|  | **Strut** | 130.88 | 129.10 | 123.90 | 122.00 | 114.13 |
|  | **Lambda** | 146.00 | 140.13 | 116.20 | 105.50 | 86.80 |
|  | **Double Mini** | 44.90 | 42.17 | 31.40 | 26.50 | 18.70 |

## 4. Discussion

This study focused on an *in silico* assessment of four plate designs of different materials for subcondylar fracture fixation during mastication cycle. Various components of mandible, including the PDL and condylar fibrous tissue, were explicitly modelled to develop the *in-silico* model. The FE study provided key insights into the influence of plate material on the mechanics of load transfer through mandible. The maximum principal strains for all the reconstructed mandibles were well below the corresponding yield strain limit of ∼2500 με [49].

The mandibular strains in the mandibles subjected to RMOL decreased with increasing elastic modulus of plate as it provided increasing stability. This trend was also exhibited under LMOL but only for mandibles reconstructed with lambda and double mini plates. For these two plates, the presence of two anterior screws in the proximal segment likely increased the resistance to the moment generated at the fracture site, thereby improving construct stability. This is also supported by the lowest interfragmentary displacements with double mini plates under both loading conditions, while lambda plates showed comparatively lower mandibular principal stresses among the configurations. For instance, under LMOL loading, the interfragmentary displacement with double mini plates ranged between approximately 18–45 μm, whereas trapezoid and strut plates showed substantially higher displacements of approximately 68–131 μm, indicating higher effective construct stiffness of the double mini configuration. In contrast, trapezoid and strut plates, which had only one anterior screw in the proximal segment, likely provided relatively lower resistance to bending moments. Consequently, the expected increase in stability with increasing plate elastic modulus was not consistently observed in these configurations. For trapezoid and strut plates, hence, the mandibular strains increased with increasing elastic modulus of plates. This trend of increasing mandibular strains was observed due to decreasing strains in the plates with increasing elastic modulus of plates (Table 6), as the softer plates undergo more deformation than stiffer plates, and absorb higher amount of total strains leading to lower strains in the corresponding reconstructed mandibles.

The study considered ipsilateral (RMOL) and contralateral (LMOL) occlusions. Under ipsilateral occlusion (RMOL), the constrained teeth remained within the fracture plane, whereas under contralateral occlusion (LMOL), the constrained teeth on the opposite side (Figure 4) caused an out-of-plane bending moment due to reaction forces. This resulted in higher forces in screws and plates under contralateral occlusion (LMOL). Consequently, as compared to ipsilateral occlusion (RMOL), higher stresses and strains were generally observed in plates under contralateral occlusion (LMOL) (Table 5 – 6).

**Figure 4.**
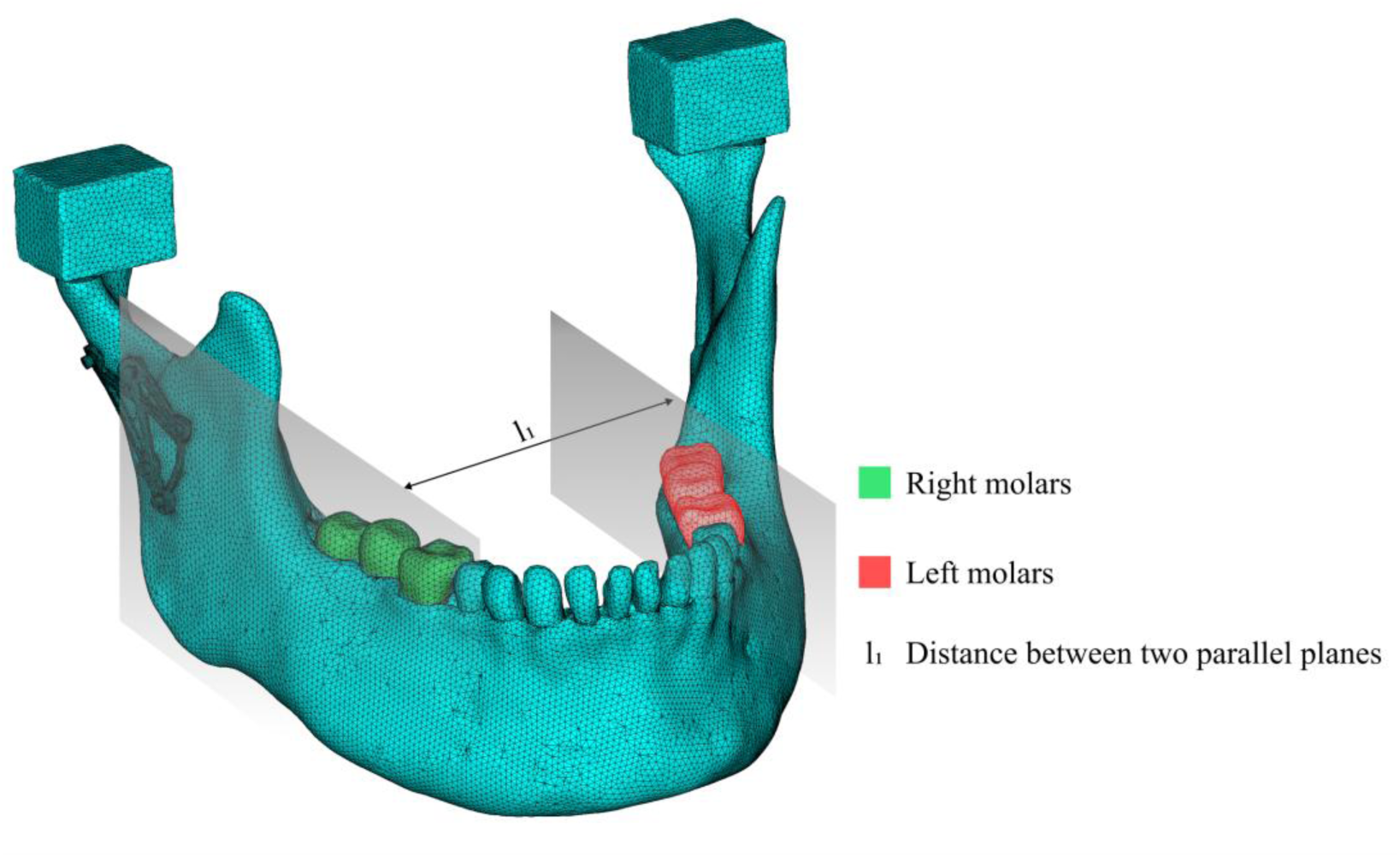
Plane passing through right molars and fracture site and plane through left molars indicating the bending effect during ipsilateral chewing forces

For further evaluation of fracture reconstruction, the possibility of failure of the reconstruction plate [50] was investigated based on the maximum von Mises stresses in the plates. As the maximum von Mises stresses in the SS316L and Magnesium plates were found to be exceeding the yield strength of 380 MPa and 162 MPa respectively [9,16], SS316L and Magnesium should be used with caution due to a risk of plate failure. With nitinol, only the use of a strut plate is suggested based on low maximum von Mises stress, supporting the favourable performance of strut plate among all designs. Both the titanium alloys (Ti-6Al-4V and TNTZ) provide adequate strength with the resulting maximum von Mises stresses well below the yield limits. TNTZ plates, even with a lower modulus than Ti-6Al-4V, had a lower ratio of maximum von Mises stress with respect to the yield strength (864 MPa [12]) than Ti-6Al-4V under LMOL for all the plate designs.

Furthermore, the stress concentration observed in the anterior arm of the trapezoid plate under LMOL can be attributed to the altered bending and torsional load transfer across the fixation system. Under contralateral occlusion, the out-of-plane bending moment generates tensile opening forces at the anterior aspect of the fracture, causing the anterior arm of the trapezoid plate to act as a cantilever resisting these forces. Additionally, the angular junction between the anterior and posterior arms of the trapezoid geometry creates a geometric discontinuity, which further promotes localized stress concentration. In contrast, under ipsilateral occlusion (RMOL), the load is applied closer to the fixation side, resulting in a shorter bending moment arm and more direct compressive load transmission through the posterior region of the plate, which might explain the marginally higher stresses observed posteriorly.

In addition to the stresses and strains, we also evaluated another important indicator of stability of fracture fixation: interfragmentary displacement. The observed reduction in the interfragmentary displacements with increase in the stiffness of the implant materials, added support to the use of stiff materials and Ti alloys. The interfragmentary displacement trend with material stiffness corroborated the results in literature [30]. However, in the design aspect, the double mini plates fixed with the highest number of screws, led to the most superior stabilisation [30]. Lambda plate resulted in a very high interfragmentary distance (beyond the critical limit of 150 μm [51,52] for facilitating fracture healing) under RMOL, indicating the risk of non-union while using lambda plate. The strut plate and trapezoid plates had almost similar interfragmentary displacements under RMOL and the interfragmentary displacements did not exceed 150 μm in any of the cases. However, the double mini plates fixed with the highest number of screws, resulted in the lowest interfragmentary displacement and least mandibular strains indicating that in terms of design, double mini design resulted in the most superior fixation. While Orassi et al. [14] showed that Mg WE43 could be used, we show that its use is configuration dependent, just like nitinol also. In terms of design, we had previously [31] found that the presence of multiple arms along the tensile and compressive strain contours led to reduced strains with trapezoid and double mini plates. We now reinforce that with a trapezoidal plate, the material selection also plays a role. Hence, this study highlights that both plate material and design strongly influence fixation stability in subcondylar fractures. Titanium alloys (Ti-6Al-4V and TNTZ) showed the most reliable performance, while SS316L and magnesium posed risks of plate failure. Among designs, double mini plates provided the most suitable stabilization with the lowest interfragmentary displacement, followed by trapezoid plates, whereas lambda plates risked non-union due to excessive displacement under ipsilateral loading. Clinically, these findings support the preferential use of Ti-based double mini or trapezoid plates [5,6] for achieving stable fixation and reliable healing, while emphasizing that material–design–loading interactions should guide patient-specific treatment planning. It should be noted that extensive clinical outcome data are currently available primarily for titanium based fixation systems [5,6], which remain the standard of care in maxillofacial fracture management. In contrast, materials such as magnesium alloys, Nitinol, and novel Ti alloys are still under active investigation, and long-term clinical evidence in subcondylar fracture fixation is limited. Therefore, the present findings for these materials should be interpreted as preclinical biomechanical insights that may guide future translational and clinical research rather than immediate clinical recommendations.

Despite the study’s thorough approach, it has a few limitations. Although heterogeneous material properties were assigned to the cancellous bone and orthotropic material properties to the cortical bone, the model does not simulate the non-linear nature of materials or the heterogeneous anisotropic properties of the cortical bone. Additionally, this study does not account for inter-patient variability owing to differences in morphological properties such as bone density and volume. Hence, the results should be considered patient specific. While biomechanical stability is a critical determinant of successful fracture fixation, clinical outcomes are also influenced by additional biological and material-related factors, such as nickel hypersensitivity and uncontrolled corrosion in magnesium-based alloys, which are not captured in the present finite element framework. However, since all comparative simulations were performed on the same anatomy under identical loading and boundary conditions, the observed differences between modeling strategies, materials, and plate designs reflect relative mechanical trends rather than subject-specific effects. Although the present study did not simulate bone healing, it is recognised that mechanical stability plays an important role in fracture repair and tissue differentiation. Recent work [53] has highlighted the mechanobiological coupling between stability and callus formation. The comparative biomechanical insights provided here thus contribute to understanding the mechanical environment that precedes biological healing. The models also use a simplified cylindrical representation of screws that likely leads to an underestimation of the local stress concentrations at the thread roots and bone-screw interface, and hence, the reported screw stress values should be interpreted as conservative lower-bound estimates.

The influence of screw placement on fixation stability is another factor that could affect the results. However, these factors were beyond the scope of the present study and planned to be investigated as an extension of this study in future. Furthermore, the present study evaluates fixation performance under static loading conditions and assesses mechanical safety primarily through stress- and strain-based metrics relative to material strength. While this approach is widely used for pre-clinical evaluation of mandibular fixation strategies, it does not explicitly capture fatigue-related failure mechanisms arising from cyclic mastication loading. Consequently, long-term failure modes such as plate fatigue or screw loosening cannot be directly inferred from the current analysis. In addition, for biodegradable materials such as the Mg WE43 alloy, progressive material degradation during the healing period may alter load-sharing behaviour and compromise fixation stability if degradation occurs faster than bone union. These time-dependent effects were beyond the scope of the present static framework and warrant future investigation.

## 5. Conclusion

A biologically realistic FE model was developed to evaluate fixation using different implant designs and materials during mastication. The different trends between plates with varying number of screws highlighted the importance of screws in the proximal segment to resist the moment present under contralateral chewing. Based on the least strains in reconstructed mandibles and the least interfragmentary distances, the double mini plates are suggested as the most optimal for reconstruction. Although, nitinol and Mg WE43 posed some advantages, they resulted in high equivalent von Mises stresses in plates, indicating the use of titanium to be more suitable. This study suggests that the patients should continue with the contralateral chewing to prevent instability. The results from this study have potential to improve development of improved fixation techniques of double mini plates of titanium alloys such as TNTZ.

## Author contributions: CRediT

Conceptualization: KM, AD; Data curation: AG, AD; Formal analysis: AG, RG, KD, AD, KM; Investigation: AG, RG, AD, KM; Methodology: AG, RG, AD, KM; Project administration: KD, KM; Resources: KD, KM; Software: KM; Supervision: KD, KM; Validation: AG, RG, AD; Visualization: AG, RG, AD, KM; Writing – original draft: AG; Writing – review and editing: AG, RG, AD, KD, KM.

Anoushka Gupta- AG, Rajdeep Ghosh-RG, Kaushik Dutta- KD, Abir Dutta- AD, Kaushik Mukherjee- KM

## Declaration of generative AI and AI-assisted technologies in the writing process

No AI tools and technologies have been used while preparation of this manuscript.

## Competing interests

None declared

## Funding

None

## Ethical approval

Not required

## Data Availability Statement

The data that support the findings of this study are available from the corresponding author upon reasonable request.

## Author Contributions Statement

This is to state that:

- all the authors were fully involved in the research design, analysis and data interpretation
- all the authors were fully involved in the preparation of the manuscript.
- all the authors also approve this final and the submitted version.

## Notes

### Competing Interest Statement

The authors have declared no competing interest.

### Summary of Updates

the manuscript has been thoroughly revised.

## References

[1] Sukegawa, S., Kanno, T., Masui, M., Sukegawa-Takahashi, Y., Kishimoto, T., Sato, A. and Furuki, Y., 2019. Which fixation methods are better between three-dimensional anatomical plate and two miniplates for the mandibular subcondylar fracture open treatment?. Journal of Cranio-Maxillofacial Surgery, 47(5), pp.771–777.

[2] Hedeșiu, M., Pavel, D.G., Almășan, O., Pavel, S.G., Hedeșiu, H. and Rafiroiu, D., 2022. Three-dimensional finite element analysis on mandibular biomechanics simulation under normal and traumatic conditions. Oral, 2(3), pp.221–237.

[3] Abdel-Galil, K. and Loukota, R., 2010. Fractures of the mandibular condyle: evidence base and current concepts of management. British Journal of Oral and Maxillofacial Surgery, 48(7), pp.520–526.

[4] Nasser, M., Pandis, N., Fleming, P.S., Fedorowicz, Z., Ellis, E. and Ali, K., 2013. Interventions for the management of mandibular fractures. Cochrane database of systematic reviews, (7).

[5] Murakami, K., Yamamoto, K., Sugiura, T., Horita, S., Matsusue, Y. and Kirita, T., 2017. Computed tomography–based 3-dimensional finite element analyses of various types of plates placed for a virtually reduced unilateral condylar fracture of the mandible of a patient. Journal of Oral and Maxillofacial Surgery, 75(6), pp.1239–e1.

[6] Wagner, A., Krach, W., Schicho, K., Undt, G., Ploder, O. and Ewers, R., 2002. A 3-dimensional finite-element analysis investigating the biomechanical behavior of the mandible and plate osteosynthesis in cases of fractures of the condylar process. Oral Surgery, Oral Medicine, Oral Pathology, Oral Radiology, and Endodontology, 94(6), pp.678–686.

[7] Anand, N. and Pal, K., 2022. Evaluation of biodegradable Zn–1Mg–1Mn and Zn–1Mg–1Mn-1HA composites with a polymer-ceramics coating of PLA/HA/TiO2 for orthopaedic applications. Journal of the Mechanical Behavior of Biomedical Materials, 136, p.105470.

[8] Prasadh, S., Ratheesh, V., Manakari, V., Parande, G., Gupta, M. and Wong, R., 2019. The potential of magnesium based materials in mandibular reconstruction. Metals, 9(3), p.302.

[9] Xie, F., He, X., Cao, S. and Qu, X., 2013. Structural and mechanical characteristics of porous 316L stainless steel fabricated by indirect selective laser sintering. Journal of Materials Processing Technology, 213(6), pp.838–843.

[10] Jacobs, J.J., Gilbert, J.L. and Urban, R.M., 1998. Current concepts review-corrosion of metal orthopaedic implants. Jbjs, 80(2), pp.268–82.

[11] Sumitomo, N., 2007. Experiment Study on Fracture Fixation with Low Rigidity Titanium Alloy-Plate Fixation of Tibia Fracture Model in Rabbit. In Proc. 21st European Conference on Biomaterials, Brighton, UK, 2007.

[12] Niinomi, M., 1998. Mechanical properties of biomedical titanium alloys. Materials Science and Engineering: A, 243(1-2), pp.231–236.

[13] Dutta, A., Mukherjee, K., Dhara, S. and Gupta, S., 2019. Design of porous titanium scaffold for complete mandibular reconstruction: The influence of pore architecture parameters. Computers in Biology and Medicine, 108, pp.31–41.

[14] Chmielewska, A. and Dean, D., 2024. The role of stiffness-matching in avoiding stress shielding-induced bone loss and stress concentration-induced skeletal reconstruction device failure. Acta Biomaterialia, 173, pp.51–65.

[15] Ghosh, R., Chanda, S. and Chakraborty, D., 2022. Application of finite element analysis to tissue differentiation and bone remodelling approaches and their use in design optimization of orthopaedic implants: A review. International Journal for Numerical Methods in Biomedical Engineering, 38(10), e3637.

[16] Orassi, V., Fischer, H., Duda, G.N., Heiland, M., Checa, S. and Rendenbach, C., 2022. In silico biomechanical evaluation of WE43 magnesium plates for mandibular fracture fixation. Frontiers in bioengineering and biotechnology, 9, p.803103.

[17] Barzegari, M., Mei, D., Lamaka, S.V. and Geris, L., 2021. Computational modeling of degradation process of biodegradable magnesium biomaterials. Corrosion Science, 190, p.109674.

[18] Jahadakbar, A., Shayesteh Moghaddam, N., Amerinatanzi, A., Dean, D., Karaca, H.E. and Elahinia, M., 2016. Finite element simulation and additive manufacturing of stiffness-matched niti fixation hardware for mandibular reconstruction surgery. Bioengineering, 3(4), p.36.

[19] Moghaddam, N.S., Skoracki, R., Miller, M., Elahinia, M. and Dean, D., 2016. Three dimensional printing of stiffness-tuned, nitinol skeletal fixation hardware with an example of mandibular segmental defect repair. Procedia Cirp, 49, pp.45–50.

[20] Lee, P.Y., Chen, Y.N., Hu, J.J. and Chang, C.H., 2018. Comparison of mechanical stability of elastic titanium, nickel-titanium, and stainless steel nails used in the fixation of diaphyseal long bone fractures. Materials, 11(11), p.2159.

[21] Aquilina, P., Chamoli, U., Parr, W.C., Clausen, P.D. and Wroe, S., 2013. Finite element analysis of three patterns of internal fixation of fractures of the mandibular condyle. British Journal of Oral and Maxillofacial Surgery, 51(4), pp.326–331.

[22] Sulukan, E. and Gümrükçü, Z., 2024. Biomechanical effects of different miniplate use on bone and miniplate systems in multiple mandible fracture: A finite element study. Injury, 55(12), p.111983.

[23] Daqiq, O., van Minnen, B., Spijkervet, F.K.L., Wubs, F.W., Lunter, G. and Roossien, C.C., 2025. Finite element analysis of mandibular fracture fixation authenticated by 3D printed mandible mechanical testing. Scientific Reports, 15(1), p.14655.

[24] Hassani, K., Ghazi, M.A. and Khorramymehr, S., 2025. A Model for Different Internal Fixation of the Mandibular Condyle Fractures. Biomedical Materials & Devices, pp.1–11.

[25] Maintz, M., Msallem, B., de Wild, M., Seiler, D., Herrmann, S., Feiler, S., Sharma, N., Dalcanale, F., Cattin, P. and Thieringer, F.M., 2023. Parameter optimization in a finite element mandibular fracture fixation model using the design of experiments approach. Journal of the Mechanical Behavior of Biomedical Materials, 144, p.105948.

[26] Pandya, H., Patel, H., Vithalani, S., Bhavsar, B., Shah, U. and Chunawala, A., 2024. Finite Element Analysis of New Modified Three-dimensional Strut Miniplate versus Conventional Plating in Mandibular Symphysis and Angle Fractures-An In vitro Study. Annals of Maxillofacial Surgery, 14(1), pp.71–75.

[27] Tazh, T., Khorramymehr, S., Hassani, K. and Nikkhoo, M., 2025. Finite element analysis of various fixation patterns in mandible bone fracture. Computer Methods in Biomechanics and Biomedical Engineering, pp.1–16.

[28] Daqiq, O., Gareb, B., Spijkervet, F.K.L., Wubs, F.W., Roossien, C.C. and van Minnen, B., 2025. Finite element analysis of the human mandible: a systematic review with meta-analysis of the essential input parameters. Scientific Reports, 15(1), p.19582.

[29] Prasadh, S., Krishnan, A.V., Lim, C.Y.H., Gupta, M. and Wong, R., 2022. Titanium versus magnesium plates for unilateral mandibular angle fracture fixation: Biomechanical evaluation using 3-dimensional finite element analysis. Journal of Materials Research and Technology, 18, pp.2064–2076.

[30] Jung, B.T., Kim, W.H., Park, B., Lee, J.H., Kim, B. and Lee, J.H., 2020. Biomechanical evaluation of unilateral subcondylar fracture of the mandible on the varying materials: A finite element analysis. PLoS One, 15(10), p.e0240352.

[31] Gupta, A., Dutta, A., Dutta, K. and Mukherjee, K., 2023. Biomechanical influence of plate configurations on mandible subcondylar fracture fixation: a finite element study. Medical & Biological Engineering & Computing, 61(10), pp.2581–2591.

[32] Polgar, K., Viceconti, M. and Connor, J.J., 2001. A comparison between automatically generated linear and parabolic tetrahedra when used to mesh a human femur. Proceedings of the Institution of Mechanical Engineers, Part H: Journal of Engineering in Medicine, 215(1), pp.85–94.

[33] Dutta, A., Mukherjee, K., Seesala, V.S., Dutta, K., Paul, R.R., Dhara, S. and Gupta, S., 2020. Load transfer across a mandible during a mastication cycle: The effects of odontogenic tumour. Proceedings of the Institution of Mechanical Engineers, Part H: Journal of Engineering in Medicine, 234(5), pp.486–495.

[34] Ichim, I., Kieser, J.A. and Swain, M.V., 2007. Functional significance of strain distribution in the human mandible under masticatory load: numerical predictions. Archives of Oral Biology, 52(5), pp.465–473.

[35] Ghosh, R., Chandra, G., Verma, V., Kaur, K., Roychoudhury, A., Mukherjee, S., Chawla, A. and Mukherjee, K., 2025. Biomechanical evaluation of temporomandibular joint implants and periprosthetic bone under unilateral and bilateral clenching. Proceedings of the Institution of Mechanical Engineers, Part H: Journal of Engineering in Medicine, 239(6), pp.560–573.

[36] Meyer, C., Serhir, L. and Boutemi, P., 2006. Experimental evaluation of three osteosynthesis devices used for stabilizing condylar fractures of the mandible. Journal of Cranio-Maxillofacial Surgery, 34(3), pp.173–181.

[37] Dutta, A., Mukherjee, K., Seesala, V.S., Dutta, K., Paul, R.R., Dhara, S. and Gupta, S., 2023. Comparative evaluation of a patient-specific customised plate designs and screws for partial mandibular reconstruction. Medical Engineering & Physics, 111, p.103941.

[38] Osteonic. [cited 2022; Available from: https://osteonic.com/_brochure/Mandible%20Plating%20System.pdf.

[39] Mukherjee, K. and Gupta, S., 2016. Bone ingrowth around porous-coated acetabular implant: a three-dimensional finite element study using mechanoregulatory algorithm. Biomechanics and modeling in mechanobiology, 15(2), pp.389–403.

[40] Mukherjee, K. and Gupta, S., 2016. The effects of musculoskeletal loading regimes on numerical evaluations of acetabular component. Proceedings of the Institution of Mechanical Engineers, Part H: Journal of Engineering in Medicine, 230(10), pp.918–929.

[41] Mukherjee, K. and Gupta, S., 2017. Mechanobiological simulations of peri-acetabular bone ingrowth: a comparative analysis of cell-phenotype specific and phenomenological algorithms. Medical & biological engineering & computing, 55(3), pp.449–465.

[42] Dalstra, M., Huiskes, R., Odgaard, A.V. and Van Erning, L., 1993. Mechanical and textural properties of pelvic trabecular bone. Journal of biomechanics, 26(4-5), pp.523–535.

[43] Reina, J.M., García-Aznar, J.M., Domínguez, J. and Doblaré, M., 2007. Numerical estimation of bone density and elastic constants distribution in a human mandible. Journal of Biomechanics, 40(4), pp.828–836.

[44] Manns A. Sistema estomatognático1988.

[45] Shockey, J.S., Von Fraunhofer, J.A. and Seligson, D., 1985. A measurement of the coefficient of static friction of human long bones. Surface Technology, 25(2), pp.167–173.

[46] Chanda, S., Mukherjee, K., Gupta, S. and Pratihar, D.K., 2020. A comparative assessment of two designs of hip stem using rule-based simulation of combined osseointegration and remodelling. Proceedings of the Institution of Mechanical Engineers, Part H: Journal of Engineering in Medicine, 234(1), pp.118–128.

[47] O’Connor, C.F., Franciscus, R.G. and Holton, N.E., 2005. Bite force production capability and efficiency in Neandertals and modern humans. American Journal of Physical Anthropology: The Official Publication of the American Association of Physical Anthropologists, 127(2), pp.129–151.

[48] Korioth, T.W., Romilly, D.P. and Hannam, A.G., 1992. Three-dimensional finite element stress analysis of the dentate human mandible. American journal of physical anthropology, 88(1), pp.69–96..

[49] Pinheiro, M., Willaert, R., Khan, A., Krairi, A. and Van Paepegem, W., 2021. Biomechanical evaluation of the human mandible after temporomandibular joint replacement under different biting conditions. Scientific Reports, 11(1), p.14034.

[50] Moiduddin, K., Anwar, S., Ahmed, N., Ashfaq, M. and Al-Ahmari, A., 2017. Computer assisted design and analysis of customized porous plate for mandibular reconstruction. Irbm, 38(2), pp.78–89.

[51] Perren, S.M., 1979. Physical and biological aspects of fracture healing with special reference to internal fixation. Clinical Orthopaedics and Related Research (1976-2007), 138, pp.175–196.

[52] Søballe, K., 1993. Hydroxyapatite ceramic coating for bone implant fixation: mechanical and histological studies in dogs. Acta Orthopaedica Scandinavica, 64(sup255), pp.1–58.

[53] Mehboob, A., Barsoum, I., Mehboob, H., Ouldyerou, A. and Al-Rub, R.K., 2025. Computational modelling and optimization of porous plates for mandibular fracture fixation accounting for bone healing. Materials & Design, 254, 114060.

