## supplemental file for "Biomechanical comparison of plate materials and designs for subcondylar fracture fixation: An *in silico* assessment"

**Table S1.** Maximum principal stress (MPa) in reconstructed mandibles under RMOL and LMOL

| | | Nitinol | Mg WE43 | TNTZ | Ti-6Al-4V | SS 316L |
|---|---|---|---|---|---|---|
| **RMOL** | **Trapezoid** | 17.96 | 18.00 | 18.11 | 18.12 | 18.08 |
| | **Strut** | 17.20 | 17.19 | 17.15 | 17.14 | 17.15 |
| | **Lambda** | 16.92 | 16.90 | 16.88 | 16.94 | 16.99 |
| | **Double Mini** | 17.77 | 17.74 | 17.63 | 17.29 | 17.51 |
| **LMOL** | **Trapezoid** | 15.91 | 15.90 | 15.89 | 15.80 | 15.86 |
| | **Strut** | 14.97 | 14.95 | 14.97 | 15.04 | 15.19 |
| | **Lambda** | 15.50 | 15.45 | 15.27 | 15.16 | 14.97 |
| | **Double Mini** | 15.77 | 15.72 | 15.53 | 15.45 | 15.33 |

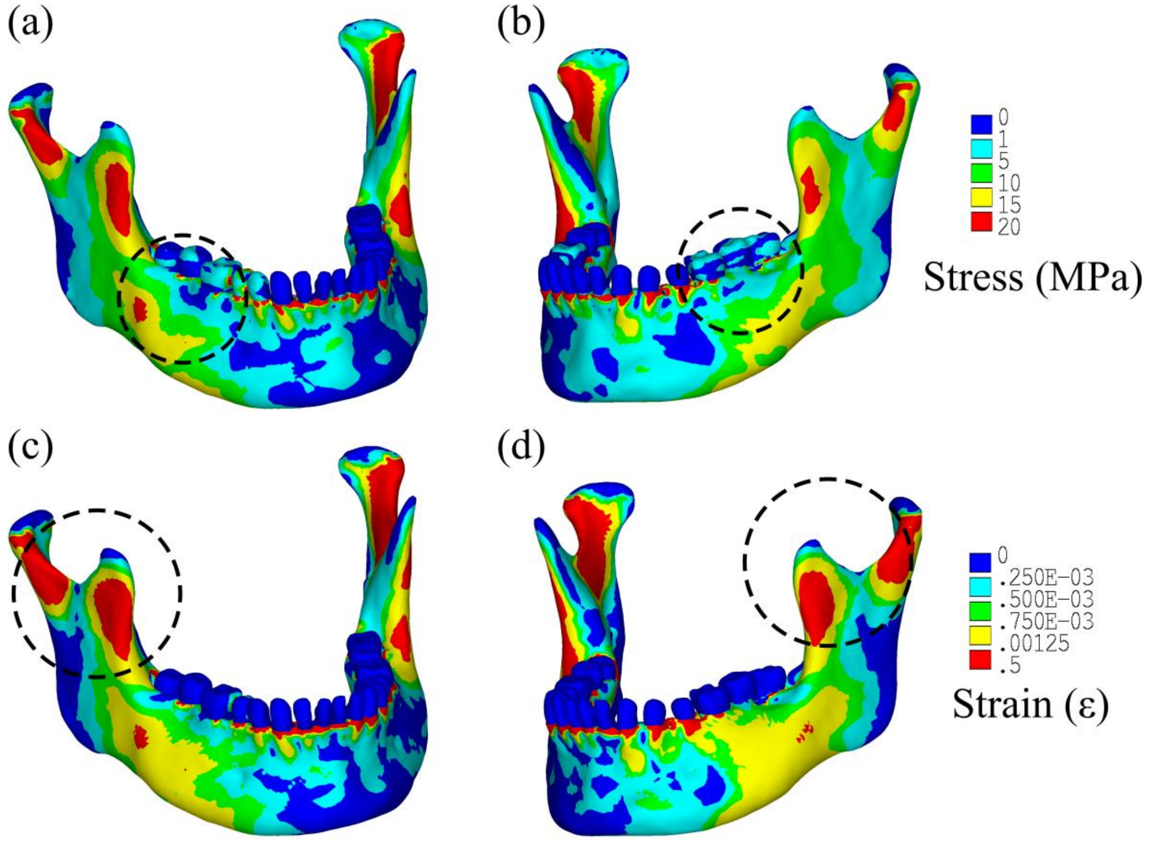


**Figure S1.** Principal tensile stress (top row) and strain (bottom row) distributions in intact mandible under (a, c) RMOL (b, d) LMOL

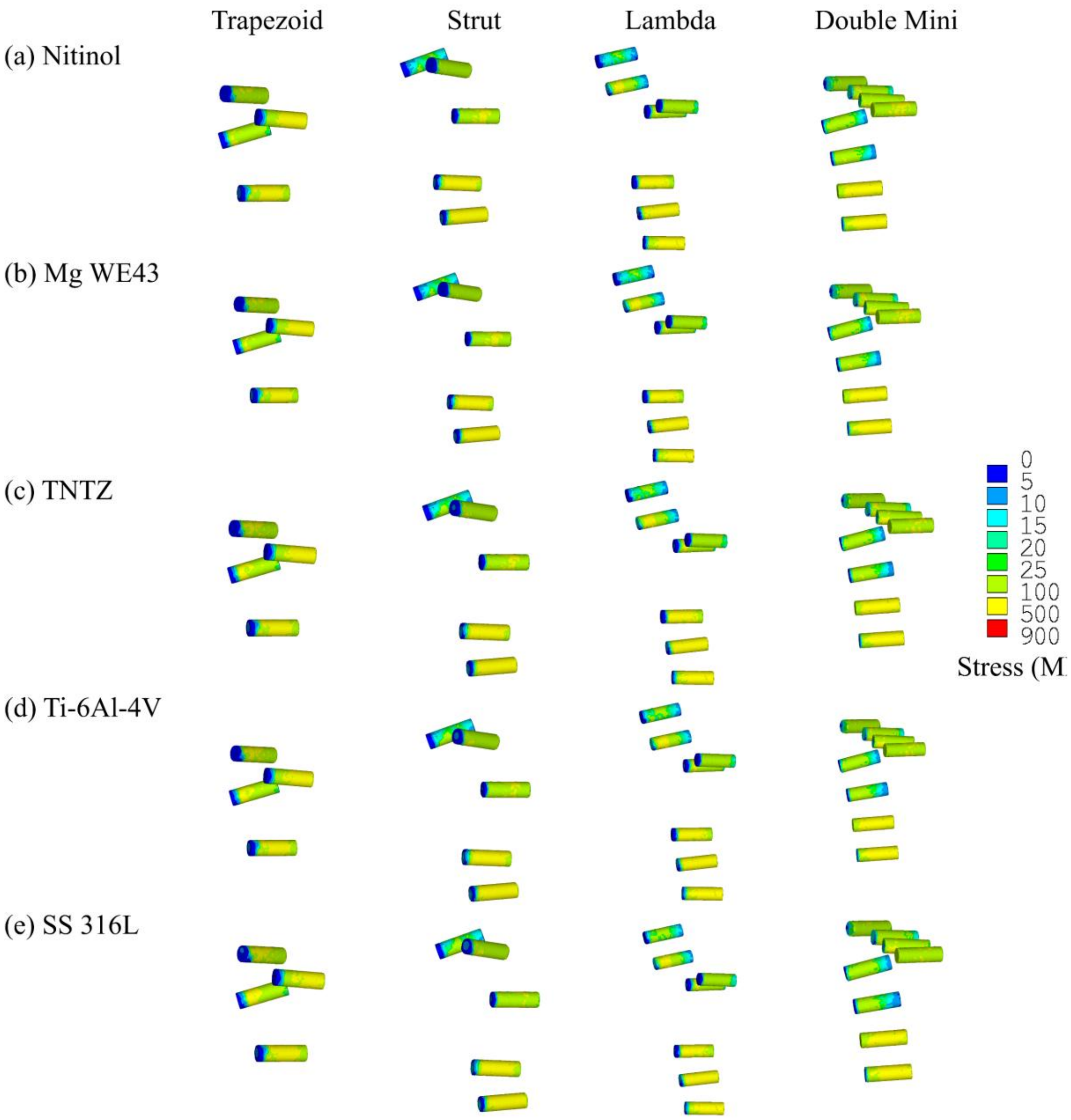


**Figure S2.** von Mises stress distribution in screws of trapezoid, strut, lambda and double mini plates for (a) Nitinol, (b) Mg WE43, (c) TNTZ, (d) Ti-6Al-4V and (e) SS 316L under RMOL

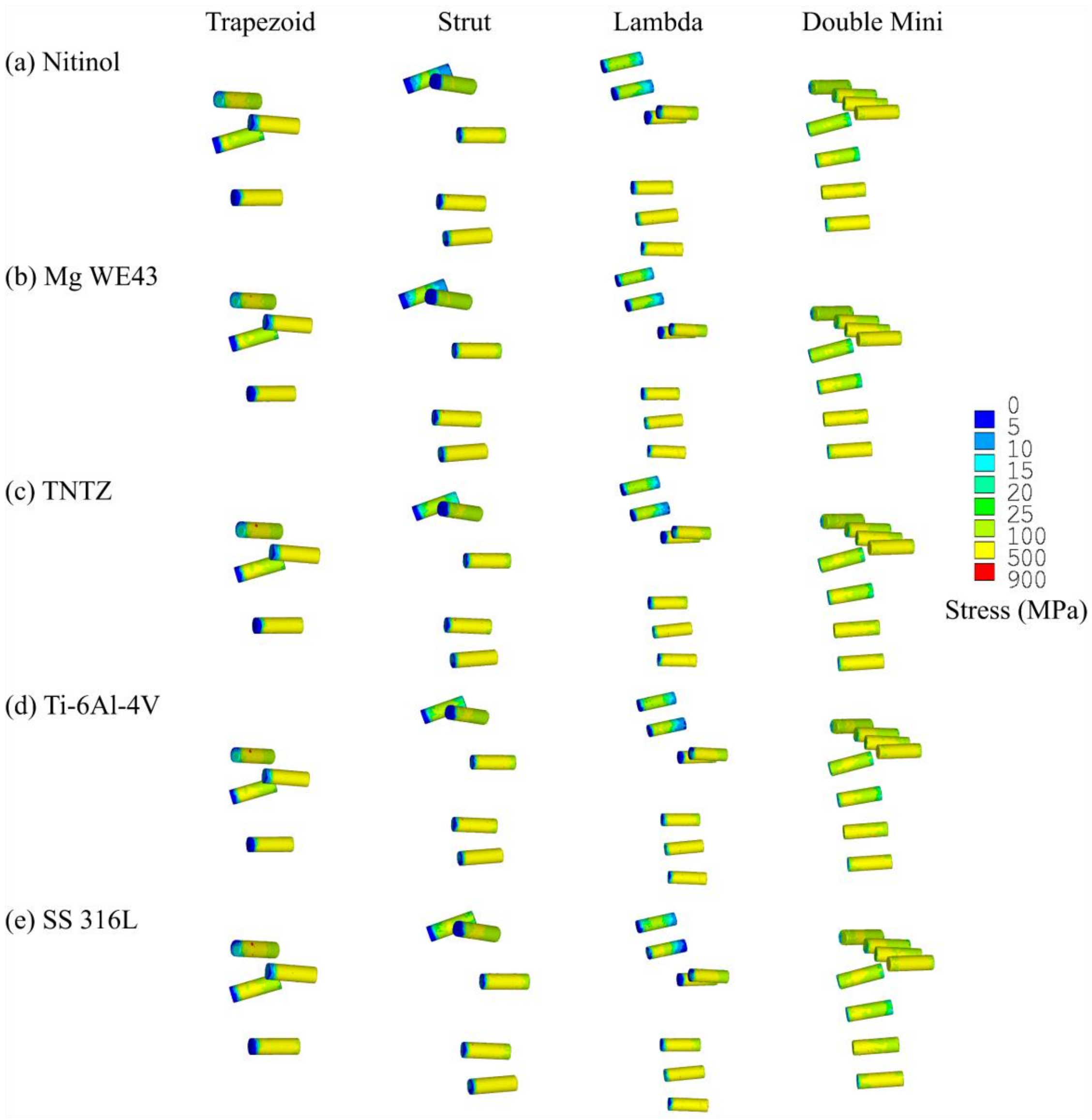


**Figure S3.** von Mises stress distribution in screws of trapezoid, strut, lambda and double mini plates for (a) Nitinol, (b) Mg WE43, (c) TNTZ, (d) Ti-6Al-4V and (e) SS 316L under LMOL
